# The exon junction complex component EIF4A3 is essential for mouse and human cortical progenitor mitosis and neurogenesis

**DOI:** 10.1101/2023.01.13.524010

**Authors:** Bianca M. Lupan, Rachel A. Solecki, Camila Manso Musso, Fernando C. Alsina, Debra L. Silver

## Abstract

Mutations in components of the exon junction complex (EJC) are associated with neurodevelopment and disease. In particular, reduced levels of the RNA helicase *EIF4A3* cause Richieri-Costa-Pereira Syndrome (RCPS) and CNVs are linked to intellectual disability. Consistent with this, *Eif4a3* haploinsufficient mice are microcephalic. Altogether, this implicates EIF4A3 in cortical development; however, the underlying mechanisms are poorly understood. Here, we use mouse and human models to demonstrate that EIF4A3 promotes cortical development by controlling progenitor mitosis, cell fate, and survival. *Eif4a3* haploinsufficiency in mice causes extensive cell death and impairs neurogenesis. Using *Eif4a3*;*p53* compound mice, we show that apoptosis is most impactful for early neurogenesis, while additional p53-independent mechanisms contribute to later stages. Live imaging of mouse and human neural progenitors reveals *Eif4a3* controls mitosis length, which influences progeny fate and viability. These phenotypes are conserved as cortical organoids derived from RCPS iPSCs exhibit aberrant neurogenesis. Finally, using rescue experiments we show that EIF4A3 controls neuron generation via the EJC. Altogether, our study demonstrates that EIF4A3 mediates neurogenesis by controlling mitosis duration and cell survival, implicating new mechanisms underlying EJC-mediated disorders.

**Summary statement:** This study shows that EIF4A3 mediates neurogenesis by controlling mitosis duration in both mouse and human neural progenitors, implicating new mechanisms underlying neurodevelopmental disorders.

## Introduction

The neocortex is responsible for higher order processes including cognition, memory, and motor control. In mice, corticogenesis initiates at mid-gestation around embryonic day 10 (E10), when neuroepithelial progenitors divide symmetrically to expand the progenitor pool (Lodato and Arlotta, 2015; Silver, 2019). These cells then give rise to radial glial cells (RGCs), which can undergo symmetric divisions to produce RGCs, or asymmetric divisions to generate excitatory pyramidal neurons or intermediate progenitors (IPs). IPs are also a major source of excitatory neurons. These excitatory neurons are produced in an “inside out” fashion, with early-born neurons forming the deep layers (VI, V) and late-born neurons generating the superficial upper layers (IV–II/III). Radially migrating neurons use the RGC scaffold to reach their final location in the cortical plate (CP) (Silver, 2019; Vaid and Huttner, 2022). In humans, these key steps of corticogenesis are largely conserved, although there are some differences in cell behavior and composition. Disruption of these processes can result in severe neurodevelopmental disorders including microcephaly and intellectual disability (Jayaraman et al., 2018).

Previous studies implicate the RNA binding exon junction complex (EJC) in cortical development and disease. The EJC is composed of three core proteins, EIF4A3, MAGOH, and RBM8A, and is a central regulator of RNA metabolism, including nonsense-mediated decay (NMD), RNA localization, and translation **(Figure 1A)** (Le Hir et al., 2016; Mao et al., 2016; Nott et al., 2004; Palacios et al., 2004). Mutations in core EJC components cause diverse neurodevelopmental disorders (McMahon et al., 2016). *EIF4A3* and *RBM8A* copy number gains and losses are both linked to intellectual disability (Nguyen et al., 2013). *RBM8A* loss of function mutations cause thrombocytopenia with absent radii (TAR syndrome), a congenital disorder associated with severe microcephaly (Albers et al., 2012). Additionally, decreased *EIF4A3* expression underlies the developmental disorder Richieri-Costa-Pereira syndrome (RCPS), in which patients exhibit craniofacial malformations as well as microcephaly (Bertola et al., 2018; Favaro et al., 2014; Hsia et al., 2018). This is largely due to repetitive 18-20 nucleotide-long motifs in the 5’UTR of *EIF4A3*. How mutations in EJC components cause these neurodevelopmental disorders remains an important question. In particular, while neural crest cell dysfunction is implicated in the developmental etiology of RCPS (Miller et al., 2017), whether neurogenesis is impaired is unknown.

**Figure 1.**
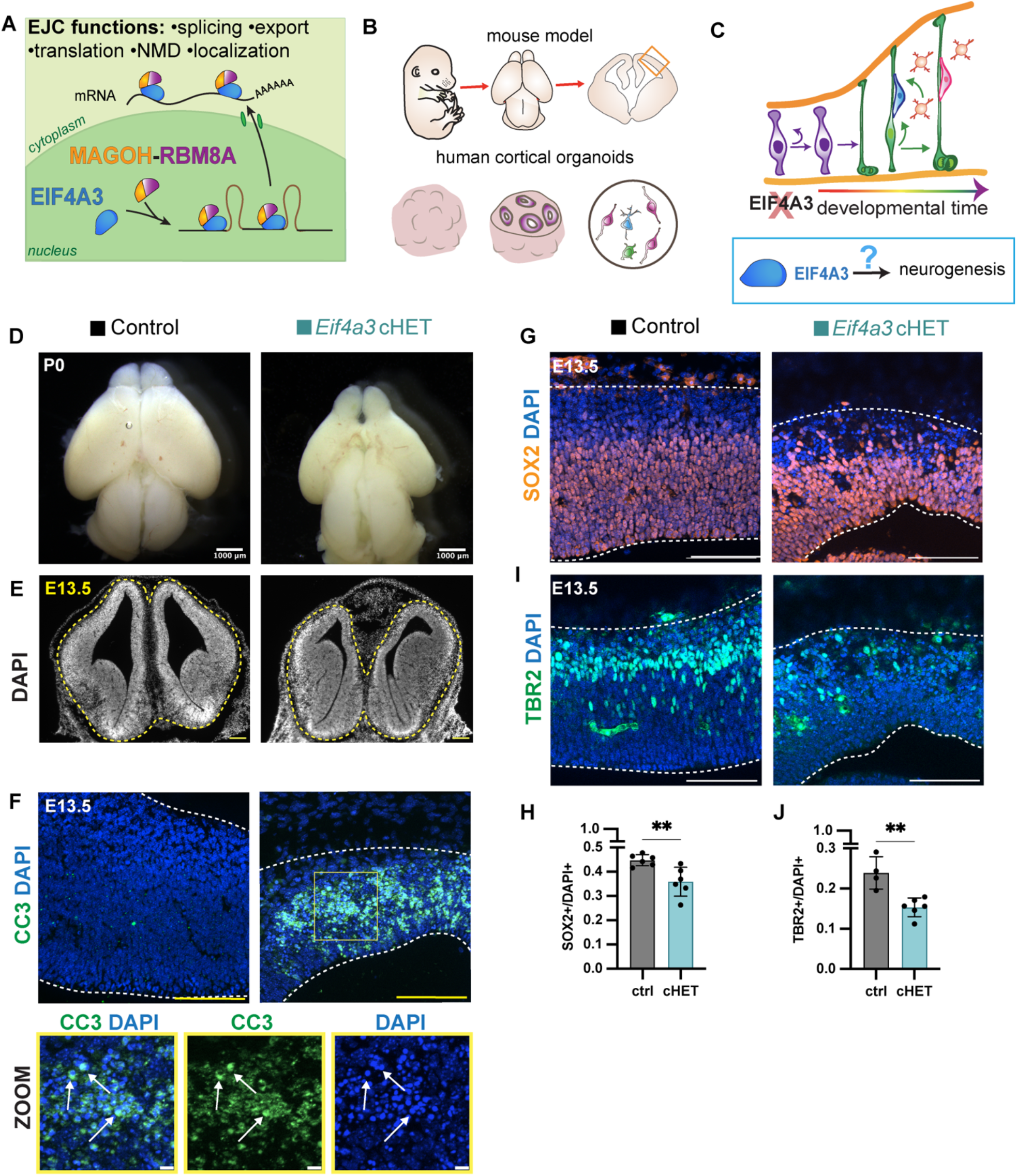
Genetic depletion of EJC component *Eif4a3* in mice causes microcephaly, massive apoptosis and loss of progenitors. **(A)** Cartoon illustrating canonical functions of the nucleo-cytoplasmic exon junction complex (EJC) in RNA metabolism. The EJC is composed of core components EIF4A3 (blue), MAGOH (orange), and RBM8A (purple) and binds mRNA at exon-exon junctions. **(B)** Top: Cartoon of a mouse embryo, embryonic brain, and coronal cortical section. Bottom: Cartoon of a human cortical organoid, a section, and a dissociated organoid culture. **(C)** Simplified cartoon of neurogenesis across developmental time. At the onset of neurogenesis, neuroepithelial progenitors (purple) undergo self-renewing divisions and then give rise to radial glia cells (RGCs, green). RGCs can undergo self-renewing divisions, generate neuron-producing intermediate progenitors (IPs, orange) or neurons (blue and pink). This study asks: what are the cellular and developmental mechanisms by which EIF4A3 regulates neurogenesis? **(D)** Images of P0 whole mount brains from indicated genotypes. **(E)** Images of E13.5 coronal sections stained for DAPI (white). Dotted yellow lines indicate borders of brain tissue. **(F)** Representative images of E13.5 coronal sections stained with DAPI (blue) and CC3 (green), with zoomed-in panels for indicated region, shown below. White arrows point to dead cells marked by CC3. **(G**,**I)** Representative images of E13.5 coronal sections stained with DAPI (blue) and **(G)** SOX2 (orange), or **(I)** TBR2 (green). Dotted lines demarcate cortex. **(H**,**J)** Quantification of **(H)** SOX2+ and **(J)** TBR2+ cells relative to all cells (DAPI) at E13.5. N=4-7 embryos per genotype. Unpaired two-tailed Student’s t-test: **P<0.01. Individual dots represent biological replicates and error bars represent s.d. Scale bars: 1000 μm (D); 200 μm (E); 100 μm (F, top; G, I), 10 μm (F, bottom).

In mouse models, haploinsufficiency for each of the core EJC components in RGCs results in microcephaly, suggesting that defective neurogenesis may contribute to these disorders (Mao et al., 2015; Mao et al., 2016; Silver et al., 2010). Transcriptomic and proteomic analyses of these embryonic brains reveal converging dysregulation of common pathways, including p53 (Mao et al., 2016). Further, *p53* genetic deletion in the *Eif4a3, Magoh*, and *Rbm8a* mutant backgrounds indicate apoptosis contributes to the microcephaly, although the mechanisms are unknown (Mao et al., 2016). Collectively these common phenotypes suggest core EJC components may work together to control neurogenesis. Previous studies of *Magoh* mutants also implicate mitosis dysregulation as a basis of microcephaly, but whether this is the case for all EJC components is unknown. This is of interest given that all 3 core components are essential for mitosis in immortalized cells (Silver et al., 2010).

In this study, we utilize mouse models as well as human cortical organoids to understand how reduced levels of *Eif4a3*, the helicase component of the EJC, influence neurogenesis and cause RCPS **(Figures 1B,C)**. Using live imaging of mouse and human progenitors, we discover conserved roles for *Eif4a3* in mitosis, cell fate, and cell death. We find that ablation of *p53* in the *Eif4a3* haploinsufficient mutant background rescues deep-layer neuron number, but not upper-layer neurons, suggesting both *p53*-dependent and -independent pathways governing progenitor behavior and cell composition. With rescue experiments, we establish EJC-dependent roles of EIF4A3 in neurogenesis. Our study reveals essential roles for cortical progenitor mitosis duration in *EIF4A3*-mediated neurodevelopmental pathologies.

## Results

### *Eif4a3* haploinsufficiency in mice causes defects in neurogenesis

We previously showed that *Eif4a3* haploinsufficiency led to microcephaly and altered progenitor and neuron number at early stages of neurogenesis (E11.5 and E12.5). To further understand how EIF4A3 controls neurogenesis and contributes to microcephaly, we employed the previously-generated *Eif4a3*^lox/+^ mice (Mao et al., 2016) and crossed it to *Emx1*-Cre. This strategy removes one copy of *Eif4a3* from RGCs and their progeny beginning at E9.5 (Chou et al., 2009; Gorski et al., 2002), resulting in *Eif4a3* conditional heterozygous brains (*Eif4a3* cHET). We analyzed *Eif4a3* cHET brains at postnatal day 0 (P0) to assess gross cortical size at the end of neurogenesis. At P0, whole mount *Eif4a3* cHET (*Emx1*-Cre;*Eif4a3*^lox/+^) brains were severely microcephalic **(Figure 1D)**. This corroborates our previous findings at E18.5 indicating that *Eif4a3* is critical for proper brain size (Mao et al., 2016).

We next evaluated neurogenesis and cortical size focusing on mid-neurogenesis stages. The cortices of E13.5 *Eif4a3* cHET brains were markedly thinner **(Figures 1E)**. As microcephaly is frequently associated with cell death, we assessed apoptosis in the *Eif4a3* cHET brains. Immunostaining for the apoptotic marker cleaved-caspase 3 (CC3) revealed massive apoptosis in *Eif4a3* cHET cortices **(Figure 1F)**. To analyze the impact of *Eif4a3* haploinsufficiency upon neurogenesis, we quantified RGC and IP populations. E13.5 *Eif4a3* cHET brains showed a 20% reduction in the density of RGCs (SOX2+/DAPI) compared to control brains **(Figures 1G,H)**. Moreover, the density of IPs (TBR2+/DAPI) was reduced by 36% **(Figures 1I,J)**. These findings corroborate previous observations of apoptosis and neurogenesis defects in younger *Eif4a3* haploinsufficient brains (Mao et al., 2016). Taken together, these analyses indicate that *Eif4a3* controls progenitor number and viability of cortical cells during different stages of neurogenesis.

### *Eif4a3* haploinsufficient progenitors are delayed in mitosis and undergo fewer viable divisions

We next sought to understand how EIF4A3 controls progenitor number. *EIF4A3, MAGOH* and *RBM8A* are each essential for mitosis in immortalized cells and EJC cHET E11.5 brains each have increased mitotic indexes (Mao et al., 2015; Mao et al., 2016; Silver et al., 2010). Additionally, *Magoh* haploinsufficiency prolongs progenitor mitosis, correlated with altered neurogenesis (Pilaz et al., 2016; Sheehan et al., 2020). Moreover, pharmacology studies indicate a causal link between this prolonged mitosis and altered cell fate in the developing cortex (Mitchell-Dick et al., 2019; Pilaz et al., 2016). Given this, we postulated that EIF4A3 may likewise influence progenitor number by affecting mitosis.

To directly test this possibility, we employed a live imaging paradigm established in our lab (Mitchell-Dick et al., 2019; Pilaz et al., 2016; Sheehan et al., 2020) **(Figure 2A)**. Primary progenitor cultures were generated from control and *Eif4a3* cHET cortices at E12.5 (Pilaz et al., 2016), a stage when neurogenesis and apoptosis defects are evident (Mao et al., 2016). Live imaging was performed for 20 hours, with images captured every 10 minutes. The average mitosis duration in *Eif4a3* cHET progenitors was 1.8-fold longer than control, taking 47 min on average compared to 26 min **(Figures 2B,C)**. This finding is consistent with the higher mitotic index seen in *Eif4a3* cHET brains (Mao et al., 2016) and the 1.8-fold longer mitoses of *Magoh* haploinsufficient progenitors relative to control at the same stage (Pilaz et al., 2016). Taken together, these analyses demonstrate that *Eif4a3* haploinsufficient progenitors exhibit prolonged mitosis *in vitro*, in addition to mitotic defects *in vivo* (Mao et al., 2016), similar to *Magoh* mutants (Pilaz et al., 2016; Silver et al., 2010).

**Figure 2.**
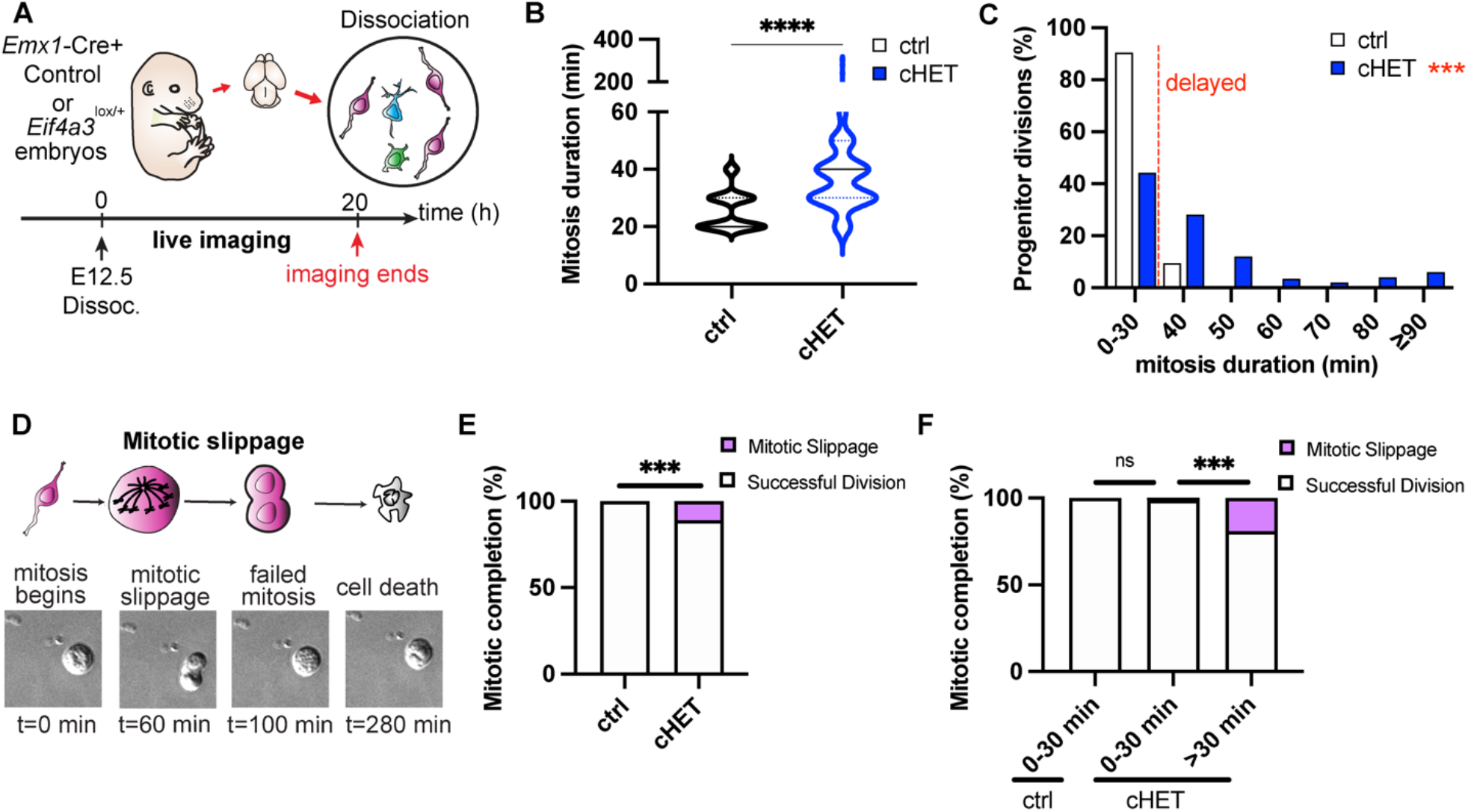
Live-imaging reveals *Eif4a3* haploinsufficient progenitors are delayed in mitosis and undergo fewer viable divisions. **(A)** Live imaging paradigm for monitoring fate of progenitors. Figure 2A was adapted from Figure 3A from Pilaz et al., 2016. **(B)** Quantification of average mitosis duration for control (black) and *Eif4a3* cHET (blue) progenitors. **(C)** Distribution of mitosis duration of control (black) and *Eif4a3* cHET (blue) progenitors. **(D)** Schematic illustrating mitotic slippage with live imaging DIC snapshots shown below, at indicated t=minutes. **(E**,**F)** Quantification of the proportion of successful divisions (white) and mitotic slippage (purple) amongst **(E)** all progeny or **(F)** relative to mitosis duration. Non-viable divisions characterized as mitotic slippage were due to unsuccessful mitosis and subsequent cell death. Unpaired two-tailed Student’s t-test: p<0.0001, **** **(B)**. χ2 analysis with post-hoc Bonferroni adjusted P-values represented by asterisks. ***P<0.001, ****P<0.0001, ns, not significant **(C**,**E**,**F)**. Error bars represent s.d. Experiments represent two live-imaging sessions, two litters; control n=3 embryos (126 cells), cHET n=3 embryos (199 cells).

We next quantified the ability of *Eif4a3* cHET progenitors to complete mitosis. We used the term “successful division” to define progenitor divisions that successfully generated two daughter cells, and the term “mitotic slippage” for divisions in which progenitors underwent a prolonged mitosis and then either senesced or died (also termed mitotic catastrophe) **(Figure 2D**) (Blagosklonny, 2007; Castedo et al., 2004). In comparison to control, 89% of *Eif4a3* cHET progenitors successfully completed mitosis, with the remainder showing mitotic slippage **(Figure 2E)**. Given the link between mitotic delay and mitotic slippage (Sheehan et al., 2020), we next asked whether progenitors undergoing slippage were mitotically delayed **(Figure 2C)**. The vast majority of control progenitors completed mitosis in 30 minutes or less, and were hence classified as “normal division” **(Figure 2C)**. Mitotically delayed *Eif4a3* cHET progenitors were significantly more likely to undergo mitotic slippage, with 19% of delayed cHET progenitors failing to complete mitosis compared to both the non-delayed cHET (1%) and control progenitors (0%) **(Figure 2F)**. Altogether, these findings indicate that *Eif4a3* controls progenitor mitosis duration and that mitotically delayed progenitors are more likely to die prior to division.

### Mitotically delayed *Eif4a3* haploinsufficient progenitors generate fewer progenitors, more neurons, and apoptotic progeny

The vast majority of *Eif4a3* cHET progenitors successfully completed mitosis. Given the neurogenesis defects observed *in vivo* **(Figure 1)** and links between mitosis duration and altered cell fate (Mitchell-Dick et al., 2019; Pilaz et al., 2016; Sheehan et al., 2020), we assessed whether *Eif4a3* cHET progenitors delayed in mitosis produced altered progeny. Towards this, we quantified viability of newborn progeny after live imaging by assaying their morphology using DIC microscopy **(Figure 3A)**. Overall, *Eif4a3* cHET progenitors produced significantly more apoptotic progeny than control progenitors **(Figures 3B,C)**, and this significantly correlated with mitotic delay **(Figure 3D)**. These data show that *Eif4a3* is required for the generation of viable progeny and that this correlates with mitotic duration.

**Figure 3.**
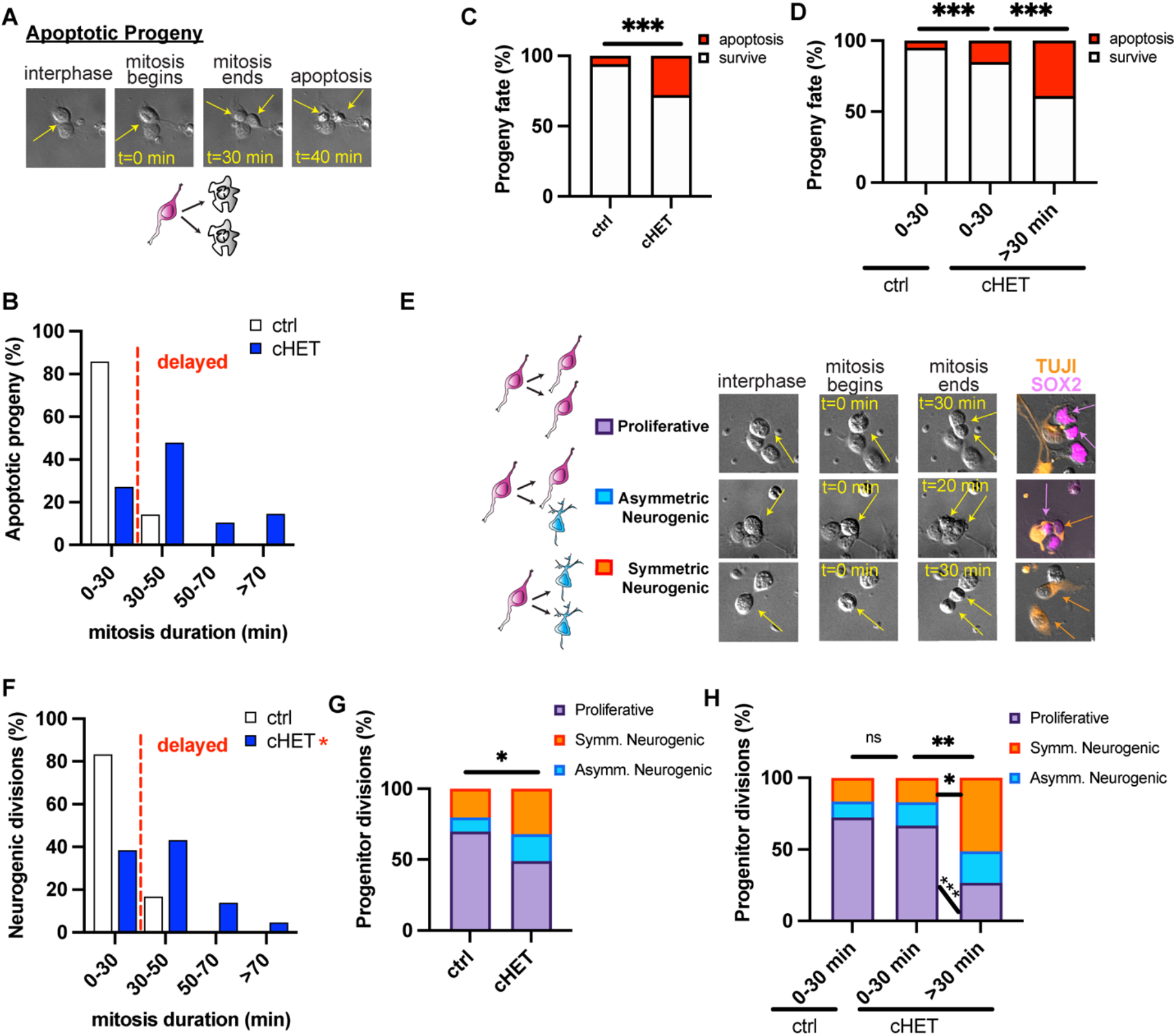
Mitotically delayed *Eif4a3* haploinsufficient progenitors produce apoptotic progeny, more neurons, and fewer progenitors. **(A)** Live imaging DIC snapshots at indicated t=minutes depicting a dividing progenitor that produces apoptotic progeny, with cartoon representation below. **(B)** Distribution of apoptotic progeny relative to mitosis duration for control (white) and *Eif4a3* cHET (blue) progenitors. **(C**,**D)** Quantification of the proportion of progenitor progeny surviving (white) or undergoing apoptosis (red) amongst **(C)** all progeny or **(D)** relative to mitosis duration. **(E)** Schematic illustrating proliferative, asymmetric neurogenic, and symmetric neurogenic divisions with examples of live and fixed analysis of progeny cell fate (right). **(F)** Distribution of neurogenic divisions relative to mitosis duration for control (white) and *Eif4a3* cHET (blue) progenitors. **(G)** Quantification of the proportion of proliferative (purple), asymmetric neurogenic (blue), or symmetric neurogenic (orange) divisions amongst **(G)** all progeny or **(H)** relative to mitosis duration. χ^2^ analysis with post-hoc Bonferroni adjusted P-values represented by asterisks; *P<0.05, **P<0.01, ***P<0.001, ns=not significant. Error bars represent s.d. Experiments represent two live-imaging sessions, two litters; control n=3 embryos (126 cells), cHET n=3 embryos (199 cells).

We next assessed the extent to which *Eif4a3* haploinsufficiency impacts the fate of surviving progeny and if there is a link to mitotic duration. Towards this, fixed progeny were stained after live imaging to identify progenitor divisions as either proliferative (SOX2+ and SOX2+ progeny), asymmetric neurogenic (SOX2+ and TUJ1+ progeny; TBR2+ and TUJ1+ progeny), or symmetric neurogenic (both TUJ1+ progeny) divisions **(Figure 3E)**. In comparison to E12.5 control and non-delayed *Eif4a3* cHET progenitors, delayed *Eif4a3* cHET progenitors underwent significantly more neurogenic divisions **(Figures 3F,G,H)**, phenocopying *Magoh*^+/−^ mutant progenitors or progenitors treated with mitotic inhibitors (Mitchell-Dick et al., 2019; Pilaz et al., 2016). Taken together, these data demonstrate that longer mitosis duration in *Eif4a3* haploinsufficient progenitors is associated with alterations in progeny cell fate and viability.

### p53-dependent and independent mechanisms explain *Eif4a3* cHET microcephaly and neurogenesis

Our data collectively indicate that *Eif4a3* controls progenitor mitosis, which is associated with altered composition of progenitors and neurons and significant apoptosis (Mao et al., 2016). A key question is the extent to which cell composition and cortical thickness changes in *Eif4a3* haploinsufficient brains are explained by apoptosis. To disentangle cell fate and cell death, we crossed *Emx1*-Cre;*Eif4a3*^lox/+^ (Mao et al., 2016) mice onto a *p53*^*lox/lox*^ background which eliminates apoptosis (Marino et al., 2000). Indeed, apoptosis was abolished in *p53*^*lox/lox*^ (p53 cKO) P0 brains as evidenced by absence of CC3+ apoptotic cells **(Figures S1A,B)**. There were no significant differences in cortical size between *Emx1*-Cre;*p53*^lox/lox^ cKO and control brains (either *Emx1*-Cre; *Eif4a3*^+/+^ or *Emx1*-Cre; *Eif4a3*^+/+^*;p53*^lox/+^) **(Figures S1D,E**). This indicates that *P53* loss alone does not grossly affect cortical size. In contrast, as expected, *Eif4a3* cHET P0 brains were strikingly reduced in size, measuring on average 55% smaller than the *Emx1*-Cre controls and missing most of the pallium **(Figures 4A,B)**. Cortical thickness in cHET brains was also significantly reduced by 54% relative to controls **(Figure 4C)**. In the *p53*;*Eif4a3* compound mutant (*Emx1*-Cre;*Eif4a3*^lox/+^*;p53*^lox/lox^), apoptosis was rescued **(Figures S1A-C)**. However, neither cortical area nor thickness were fully restored in the compound mutant, suggesting that additional factors independent of *p53* and apoptosis contribute to cortical size **(Figures 4A-C)**.

**Figure 4.**
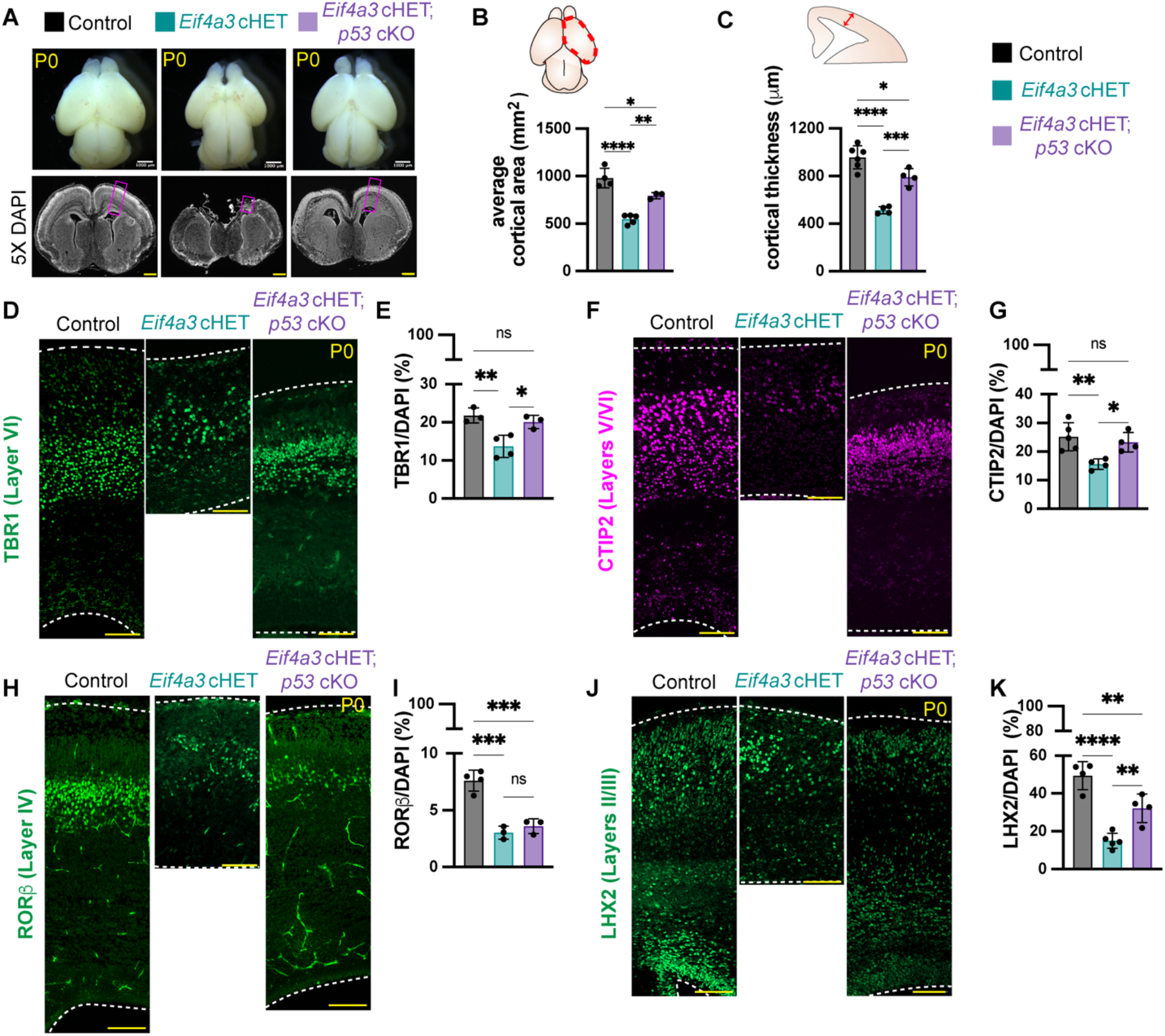
Loss of *p53* partially rescues microcephaly and neuron number associated with *Eif4a3* haploinsufficiency. **(A)** Images of P0 whole mount brains from indicated genotypes (top). Low magnification images of P0 coronal sections, stained for DAPI (white). Purple boxes denote region of coronal sections that were stained in (**D,F,H,J). B)** Quantification of cortical area and **(C)** cortical thickness in P0 brains with indicated genotypes. **(D,F,H,J)** Representative sections at P0 of indicated genotypes stained with **(D)** TBR1 (green), **(F)** CTIP2 (magenta), **(H)** ROR (green), and **(J)** LHX2 (green). **(E,G,I,K)** Quantification of **(E)** TBR1+, **(G)** CTIP2+, **(I)** ROR+, **(K)** LHX2+ cell density at P0. Dotted lines demarcate dorsal cortex. N=3-7 embryos per genotype for **(A-K)**. ANOVA with Tukey post-hoc: *P<0.05, **P<0.01, ***P<0.001, ****P<0.001, ns=not significant. Individual dots represent biological replicates and error bars represent s.d. Scale bars: 1000 μm (**A top**), 500 μm (**A bottom**); 100 μm (**D,F,H,J**).

We next used this genetic paradigm to assess the extent to which apoptosis explains altered neuronal composition of *Eif4a3* haploinsufficient brains. Toward this, we first quantified deep-layer early-born neurons (Greig et al., 2013) **(Figures 4D-K)**. In the *Eif4a3* cHET brains, immunostaining revealed 37% fewer TBR1+ (Layer VI) and 45% fewer CTIP2+ (Layer V/VI) deep-layer neurons compared to control **(Figures 4D-G)**. Notably, TBR1 and CTIP2 neuron populations were both significantly rescued in the *p53*;*Eif4a3* compound mutant brains **(Figures 4D-G)**. These data therefore suggest that reduced numbers of deep-layer neurons are largely explained by *p53* activation and apoptosis.

To examine later-born superficial layer neurons (Greig et al., 2013), we performed immunostaining for ROR (Layer IV) and LHX2 (Layer II/III) **(Figures 4H-K)**. Compared to control, *Eif4a3* cHET brains had 61% fewer ROR+ neurons while *p53*;*Eif4a3* compound mutant brains still had 53% fewer ROR+ neurons, indicating these neurons were not significantly rescued **(Figures 4H-K)**. Similarly, LHX2+ neurons were markedly reduced by 70% in *Eif4a3* cHET brains compared to control, but only partially rescued in the *p53*;*Eif4a3* compound mutant brains which had 41% fewer LHX2+ neurons compared to control **(Figures 4J-K)**. Thus, both LHX2 and ROR neurons are partially, but not completely, rescued in the *p53* mutant background. This indicates that loss of superficial neurons in *Eif4a3* mutants is due in part to p53-mediated cell death but additionally, that p53-independent mechanisms contribute to their genesis.

We next sought to understand whether differential cell death of progenitors at early and later stages of development explains the subtype-specific neuronal phenotypes at P0. The genesis of layer VI and V neurons peaks from E12.5-E13.5, and genesis of layers IV and II/III peaks from E14.5-E16.5, respectively (Greig et al., 2013). Thus, we first used our compound mutant brains to evaluate progenitors at E13.5 to understand genesis of deep-layer neurons. Staining for CC3 in the *p53;Eif4a3* compound mutant brains at E13.5 verified that apoptosis is fully rescued at this stage **(Figure S2A)**. Likewise, the cortical thickness of control brains and *p53;Eif4a3* brains was similar **(Figure 5A,B)**. The density of RGCs and IPs were also both significantly rescued in the *p53;Eif4a3* compound brains **(Figures 5C-F)**. When reported as a total number of progenitors, these trends were also present (**Figure S2B,C)**. These findings show that at E13.5, p53-mediated cell death is largely responsible for both reduced cortical thickness and reduced progenitors in *Eif4a3* haploinsufficient brains.

**Figure 5.**
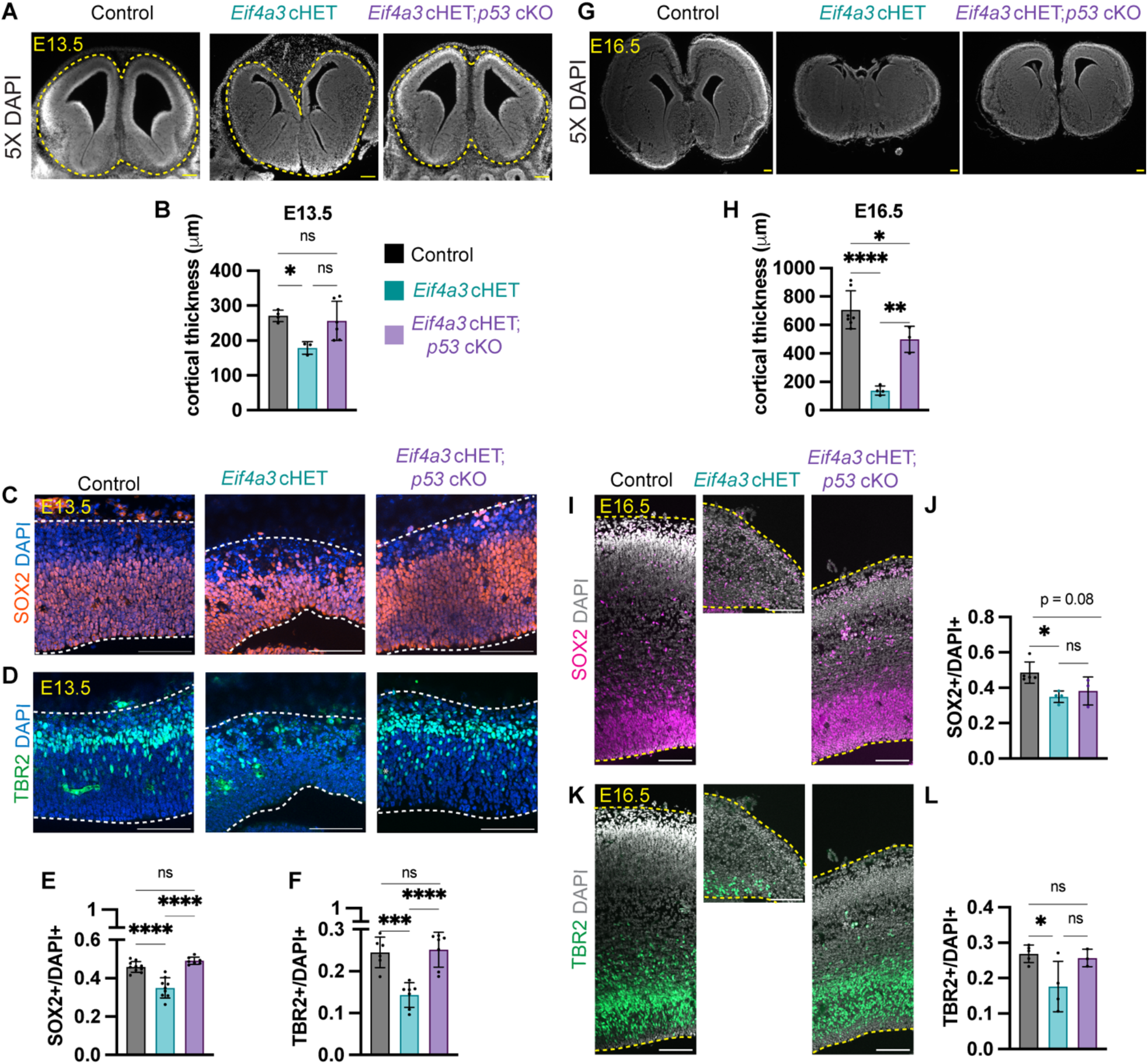
Genetic ablation of *p53* significantly rescues *Eif4a3*-dependent cortical thickness and progenitor number at E13.5, but not E16.5. **(A)** Low magnification images of coronal sections at E13.5, stained for DAPI (white). **(B)** Quantification of cortical thickness in E13.5 brains with indicated genotypes. **(C,D)** Representative images of coronal E13.5 sections stained with **(C)** SOX2 (orange) and DAPI (blue) or **(D)** TBR2 (green) and DAPI (blue). **(E,F)** Quantification of **(E)** SOX2+ cells and **(F)** TBR2+ cell density at E13.5. **(G)** Low magnification images of coronal sections at E16.5, stained for DAPI (white). **(H)** Quantification of cortical thickness in E16.5 brains with indicated genotypes. **(I,K)** Representative images of coronal E16.5 sections stained with **(I)** SOX2 (magenta) and DAPI (white) or **(K)** TBR2 (green) and DAPI (white). **(J,L)** Quantification of **(J)** SOX2+ or **(L)** TBR2+ cell density at E16.5. For E13.5: N=7-10 embryos per genotype **(A-F)**. For E16.5: N=3-5 embryos per genotype **(G-L)**. Dotted lines demarcate dorsal cortex. ANOVA with Tukey post-hoc: *P<0.05, **P<0.01, ***P<0.001, ****P<0.001, ns=not significant. Individual dots represent biological replicates and error bars represent s.d. Scale bars: 200 μm **(A)**,100 μm **(C, D, G. I, K)**.

We next examined E16.5 stages when progenitors produce layer IV and II/III neurons (Greig et al., 2013). We first measured cortical area of whole-mount brains and thickness in cortical sections. The cortex was almost entirely absent at E16.5 in the *Eif4a3* cHET brain, but this was partially (77%) restored in the *p53;Eif4a3* compound **(Figure 5G, S2D)**. Similarly, cortical thickness was 71% restored in compound mutants, compared to control **(Figure 5H)**. Thus, unlike at E13.5, at E16.5, apoptosis is insufficient to fully explain the microcephaly associated with *Eif4a3* haploinsufficiency.

We next assessed the extent to which apoptosis explains reduced RGC and IP populations in *Eif4a3* haploinsufficient brains **(Figures 5I-L)**. The density of both progenitor populations was significantly reduced in the *Eif4a3* cHET brains, consistent with E13.5 **(Figures 5E,F,J,L)**. In the *p53;Eif4a3* compound mutant brains, neither RGCs nor IPs showed significant recovery, although IPs trended **(Figures 5J,L, S2E,F)**. Thus, the reduction of upper-layer neurons at P0 is likely due to both apoptosis and additional defects in cell fate specification. Altogether, our data provide evidence of p53-dependent and p53-independent mechanisms of EIF4A3 over the course of development.

### *EIF4A3* depletion from human NPCs results in fewer proliferative and viable divisions

Having found that *Eif4a3* is essential for neurogenesis in our mouse model, we next asked whether this requirement is conserved in humans. Towards this we dissociated neural progenitor cells (NPCs) from D35 cortical organoids, electroporated these with siRNAs against *EIF4A3* or scrambled, and allowed for two days of knockdown **(Figure 6A)**. We quantified a 68% reduction in *EIF4A3* in electroporated cells after 48 hours **(Figure 6B)**. We then quantified mitosis and neurogenesis of individual progenitors. A similar live imaging paradigm as in **Figure 3** was performed except that progeny were immunostained for Ki67 (proliferative marker) and Tuj1 (neuronal marker). Notably, *EIF4A3*-deficient hNPCs were significantly delayed in mitosis compared to control siRNA-treated progenitors **(Figure 6C)**. The average mitosis duration of control NPCs was 36 minutes whereas it was 54 minutes in *EIF4A3*-deficient NPCs. To quantify a potential link between mitosis duration and progeny of these progenitors, we classified normal hNPC mitosis duration as ≤40 min, as 91.4% of control cells completed mitosis in 40 minutes or less **(Figure 6D)**. 88% of *EIF4A3*-deficient NPC divisions that produced apoptotic progeny were mitotically-delayed **(Figure 6E)**. In comparison to either control siRNA-treated NPCs or non-delayed *EIF4A3*-deficient NPCs, delayed *EIF4A3*-deficient NPCs produced significantly more apoptotic progeny **(Figures 6F,G)**. There was also an association between mitotic delay and altered neurogenic divisions. Delayed *EIF4A3*-deficient hNPCs were significantly more likely to undergo neurogenic divisions compared to non-delayed progenitors **(Figures 6H,I)**. These findings parallel the mouse data, demonstrating that *EIF4A3* controls mitosis of human neural progenitors, which is clearly linked to altered fate of progeny. Hence, the requirement of EIF4A3 for neurogenesis is evolutionarily conserved.

**Figure 6.**
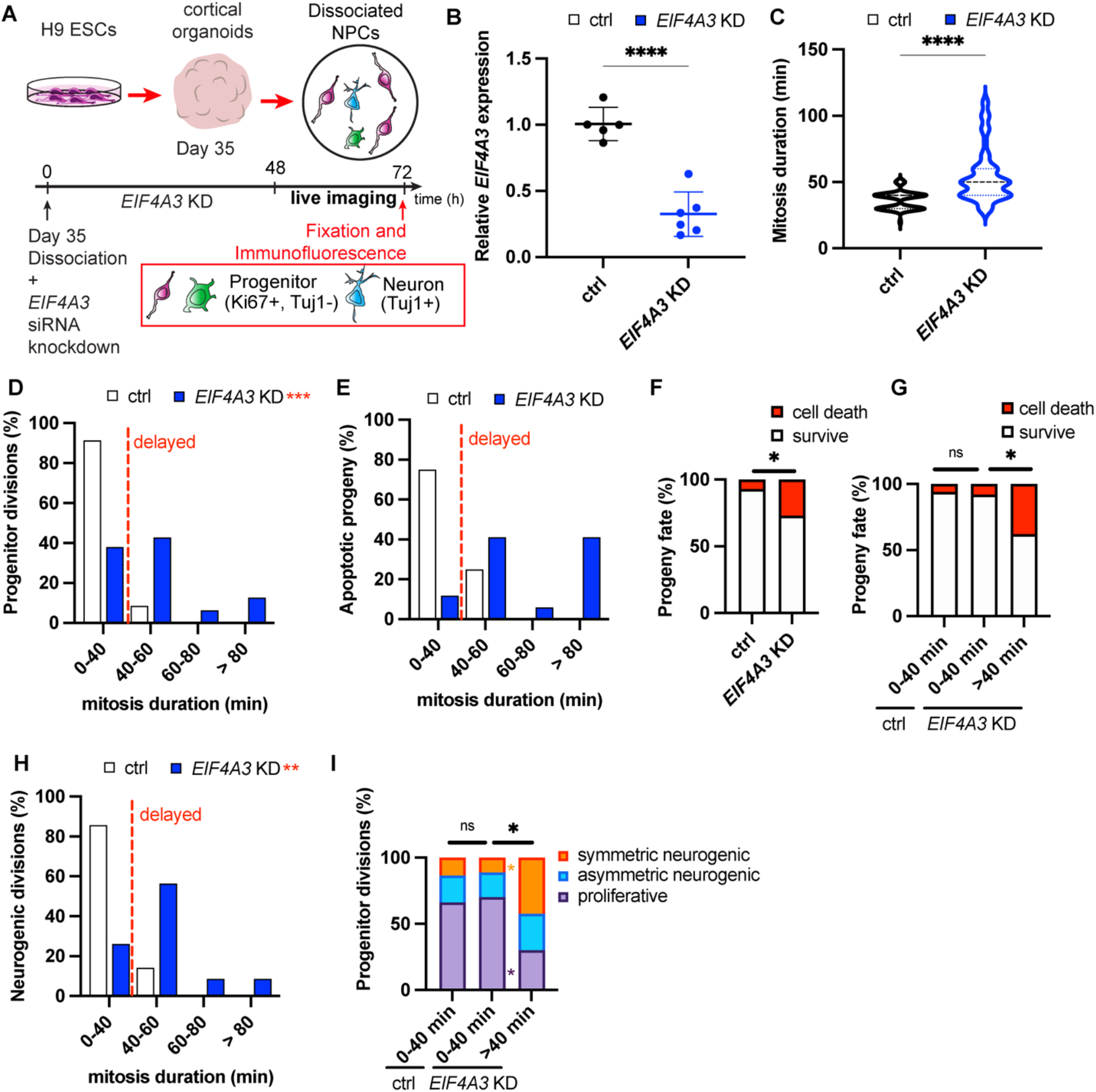
*EIF4A3* depletion from human NPCs causes mitotic delay and altered progeny fate and survival. **(A)** Schematic depicting live-imaging paradigm for human NPCs (hNPCs) dissociated from Day 35 cortical organoids. The dissociated hNPCs were electroporated with control (scrambled) or *EIF4A3* siRNAs, cultured for 48 hours, imaged for 24 hours, and subsequently stained to assign progeny cell fate. Figure 5A has been adapted from Figure 3A and E from Pilaz et al., 2016. **(B)** qPCR quantification of *EIF4A3* mRNA levels, normalized to *TBP*, in siRNA-treated hNPC cultures. **(C)** Quantification of average mitosis duration for control (black) and *EIF4A3* siRNA-treated (blue) hNPCs. **(D)** Distribution of mitosis duration of control (black) and *EIF4A3* siRNA-treated (blue) hNPCs. **(E)** Distribution of apoptotic progeny relative to mitosis duration of control (black) and *EIF4A3* siRNA-treated (blue) hNPCs. **(F,G)** Quantification of the proportion of hNPC progeny surviving (white) or undergoing apoptosis (red) amongst **(F)** all progeny or **(G)** relative to mitosis duration. **(H)** Distribution of neurogenic divisions relative to mitosis duration for control (black) and *EIF4A3* siRNA-treated (blue) hNPCs. **(I)** Quantification of the proportion of proliferative (purple), asymmetric neurogenic (blue), or symmetric neurogenic (orange) divisions relative to mitosis duration. Unpaired two-tailed Student’s t-test: p<0.0001, **** (**B,C**). χ^2^ analysis with post-hoc Bonferroni adjusted P-values. *P<0.05, **P<0.01, ***P<0.001, ****P<0.0001, ns=not significant (**D,E,F,G,H,I**). Error bars represent s.d. Experiments represent two live-imaging sessions, three independent electroporations per genotype; control n=58 cells, *EIF4A3* KD=63 cells.

### Human RCPS cortical organoids exhibit altered neurogenesis

Human *EIF4A3* hypomorphic mutations cause Richieri-Costa-Pereira syndrome (RCPS), a rare developmental disorder associated with microcephaly and language and learning disability, among other developmental deficits (Bertola et al., 2018; Favaro et al., 2014; Hsia et al., 2018). How reduced *EIF4A3* expression causes RCPS neurological defects is largely unknown. To address this gap, we utilized previously characterized patient-derived iPSCs to generate cortical organoids that model early human brain development (Alsina et al., 2022; Miller et al., 2017; Yoon et al., 2019) **(Figure 7A)**. We previously showed that in Day 35 (D35) organoids, *EIF4A3* is reduced by about 50% (Alsina et al., 2022). To assess progenitor number, we immunostained organoid sections for SOX2. RCPS organoids at D35 had 35% fewer RGCs compared to control **(Figures 7B,C)**. Immunostaining for SOX2 and PH3 (PH3+SOX2+/SOX2+) revealed on average 1.5-fold more mitotic RGCs in RCPS organoids **(Figures 7B,D)**. This demonstrates that RCPS patient-derived organoids contain fewer progenitors, but that these progenitors show a higher mitotic index.

**Figure 7.**
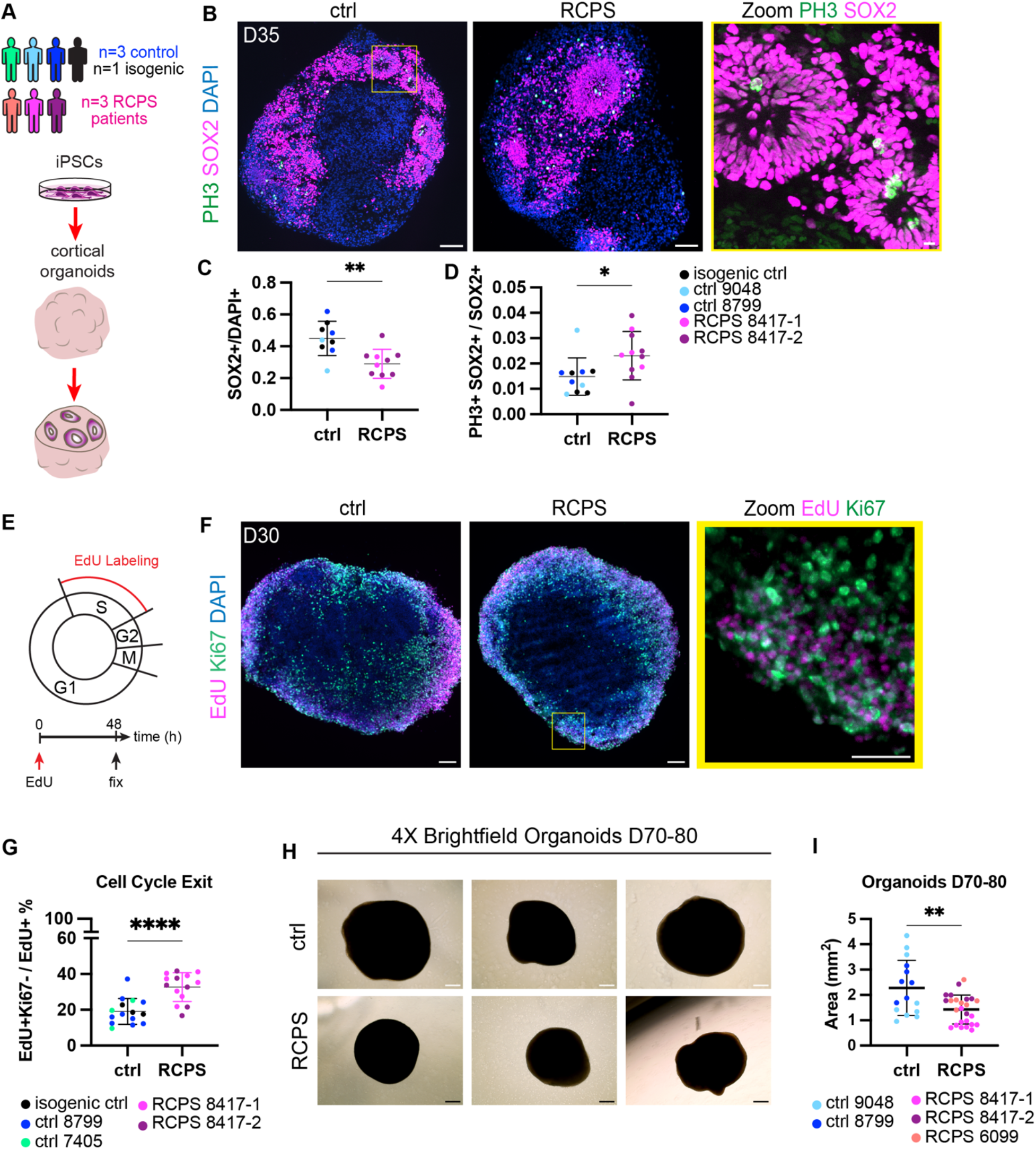
*EIF4A3* RCPS human cortical organoids exhibit neurogenesis defects. **(A)** Schematic representing the generation of cortical organoids and organoid sections from 3 control (ctrl, green, light blue, dark blue), 1 isogenic (black), and 3 Richieri-Costa-Pereira Syndrome (RCPS) (orange, pink, purple) patient-derived iPSCs. Modified from Figure 3A in Alsina et al., 2022. **(B)** Representative images of ctrl and RCPS organoids at D35 stained for SOX2 (magenta), PH3 (green), and DAPI (blue). Yellow box: higher magnification inset of PH3 and SOX2 staining. **(C)** Quantification of SOX2+ (magenta) cells relative to all cells in organoid section (DAPI). N=9 control, 10 RCPS. **(D)** Quantification of PH3+ SOX2+ (green, magenta) cells relative to all SOX2+ (magenta) cells in organoid section (DAPI). N=10 control, 11 RCPS. **(E)** Schematic illustrating the EdU labeling paradigm. **(F)** Representative images of ctrl and RCPS organoids at D30 pulsed with EdU for 2 days and stained for EdU (magenta), Ki67 (green), and DAPI (blue), Zoomed image of EdU and Ki67 staining on the right **(G)** Quantification of EdU+ Ki67+ (magenta, green) cells relative to all labeled EdU+ (magenta) cells. N=15 control, 14 RCPS. **(H)** Representative brightfield images of ctrl and RCPS organoids at D70-80. **(I)** Quantification of ctrl and RCPS organoids area at D70-80. N=15 control, 25 RCPS. Analyses were made from two or more independent differentiations. Individual dots represent one organoid (**C,D,G,I**). Unpaired two-tailed Student’s t-test: *P<0.05, **P<0.01, ***P<0.001, ****P<0.0001 (**C,D,G,I**). Scale bar: 100 μm **(B left, F left, H**). 50 μm (**F right**). 10 μm **(B right)**.

To assess neuronal generation in RCPS organoids, we quantified cell cycle exit. Towards this, we performed a 48-hour pulse of the nucleotide analog EdU which is incorporated in proliferating cells during S phase **(Figure 7E)**. Quantification of EdU and Ki67 revealed significantly more cells which have exited the cell cycle (EdU+Ki67-/EdU+) in the RCPS organoids **(Figures 7F,G)**. These data are consistent with the finding that RCPS organoids have fewer progenitors, and also indicate that *Eif4a3* regulation of cell fate is conserved in mouse and humans.

Given the microcephaly observed in some RCPS patients and in *Eif4a3* cHETs, we next asked whether these neurogenesis defects were associated with reduced size in RCPS organoids. Indeed, RPCS organoids at D70-80 were 38% smaller on average compared to control organoids (p=0.0022) **(Figures 7H,I)**. The smaller overall size of RCPS organoids fits with our data showing altered neurogenesis, further mirroring the phenotypes observed in *Eif4a3* haploinsufficient brains.

### EIF4A3 acts in an EJC-dependent manner to control newborn neuron number

EIF4A3 is a canonical regulator of RNA metabolism, as one of the three core components of the EJC. While individual EJC mutants show similar brain phenotypes and transcriptome changes (Mao et al., 2016), the extent to which EIF4A3 actually functions in this complex during neurogenesis is unknown. To address this we used a point mutant, EIF4A3^T163D^ (Ryu et al., 2019), previously shown to abolish EIF4A3 binding to the MAGOH-RBM8A heterodimer and RNA. We asked whether EIF4A3^WT^ or EIF4A3^T163D^ (Alsina et al., 2022; Ryu et al., 2019) is sufficient to rescue *Eif4a3* cHET neurogenesis phenotypes **(Figure 8A)**. Primary progenitor cultures were generated from E12.5 control and *Eif4a3* cHET cortices and electroporated with one of the following plasmid combinations: 1) GFP, 2) GFP and 3x-FLAG-EIF4A3^WT^, or 3) GFP and 3x-FLAG-EIF4A3^T163D^ **(Figures 8A,B)**. The GFP plasmid served as a readout for electroporated cells, and expression of the 3x-FLAG-EIF4A3 constructs was validated by FLAG immunostaining **(Figure S3A-F)**. After 48 hours, the primary cultures were fixed and stained for TUJI to mark neurons. For control cultures, there was no significant difference in the proportion of neurons between the GFP alone, EIF4A3^WT^, and EIF4A3^T163D^ conditions (31%, 31%, and 36% TUJ1+GFP+/GFP+, respectively) **(Figures 8C-E, I)**. Thus, with the amounts used, neither EIF4A3^WT^ nor EIF4A3^T163D^ alone caused an overexpression phenotype (see methods). For *Eif4a3* cHET cultures, there was a significant 1.8-fold increase in the proportion of GFP+ neurons (56% compared to 31% for control) **(Figure 8F-I)**. These proportions are consistent with increased neurons seen in *Eif4a3* cHET fixed brains (Mao et al., 2016) and in live imaging **(Figures 2,3)**. As predicted, expression of EIF4A3^WT^ rescued premature neurogenesis of *Eif4a3* cHET progenitors **(Figures 8G,I)**. However, expression of the EJC mutant, EIF4A3^T163D^, did not rescue these defects in *Eif4a3* cHET progenitors (59% TUJ1+GFP+/GFP+, not significant compared to GFP alone) **(Figure 8H,I)**. All together, these rescue experiments suggest the core EJC components EIF4A3, MAGOH, and RBM8A, are likely working in a complex to control neuronal generation **(Figure 9)**.

**Figure 8.**
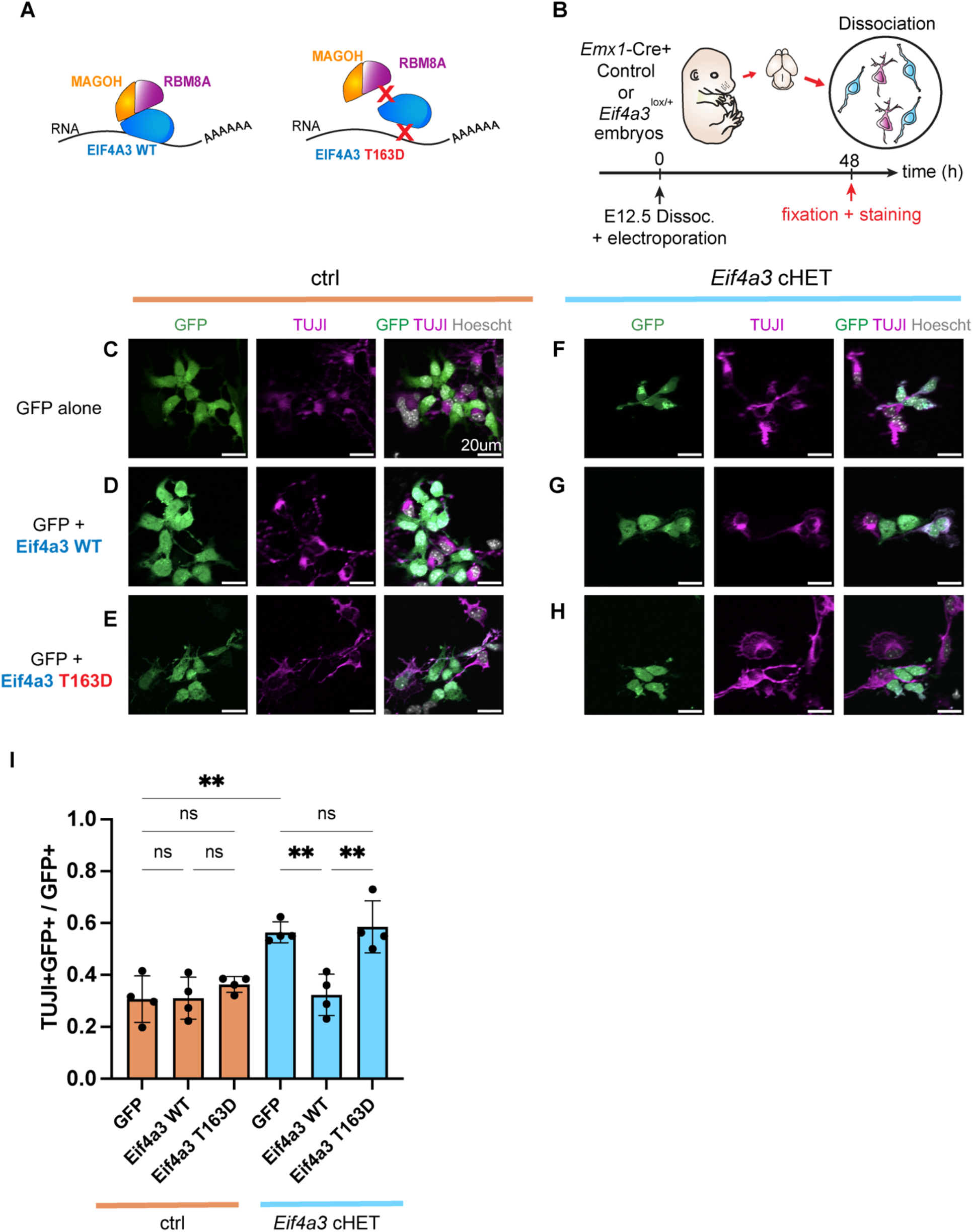
EIF4A3 acts via the EJC to control neuron generation. **(A)** Schematic depicting EIF4A3^WT^ (blue) bound to mRNA and the MAGOH-RBM8A heterodimer (orange and purple) and point mutant EIF4A3^T163D^ which is unable to bind to RNA and the MAGOH-RBM8A heterodimer. Adapted from Figure 2H-I in Alsina et al., 2022. **(B)** Schematic of rescue experiments in which E12.5 control (*Emx1*-Cre;*Eif4a3*^+/+^) or cHET (*Emx1*-Cre;*Eif4a3*^*l*ox/+^) primary cultures were electroporated with pCAG-GFP and the pCAG-3x-FLAG constructs. Cultures were fixed after 48 hours and stained for GFP (green), TUJI (magenta), and Hoescht (white); representative images are shown in **(C-H). (I)** Quantification of the proportion of electroporated neurons (GFP+, TUJI+; green, magenta) relative to the total number of electroporated cells (GFP; green). Control cells are graphed in orange and cHET are graphed in light blue, with the electroporation conditions listed on the x-axis. N=1135 ctrl cells, N=503 cHET cells, 4 embryos from 2 independent litters. ANOVA with Tukey post-hoc: *P<0.05, **P<0.01, ***P<0.001, ****P<0.001, ns=not significant. Individual dots represent biological replicates and error bars represent s.d. scale bars: 20 μm **(C-H)**.

**Figure 9.**
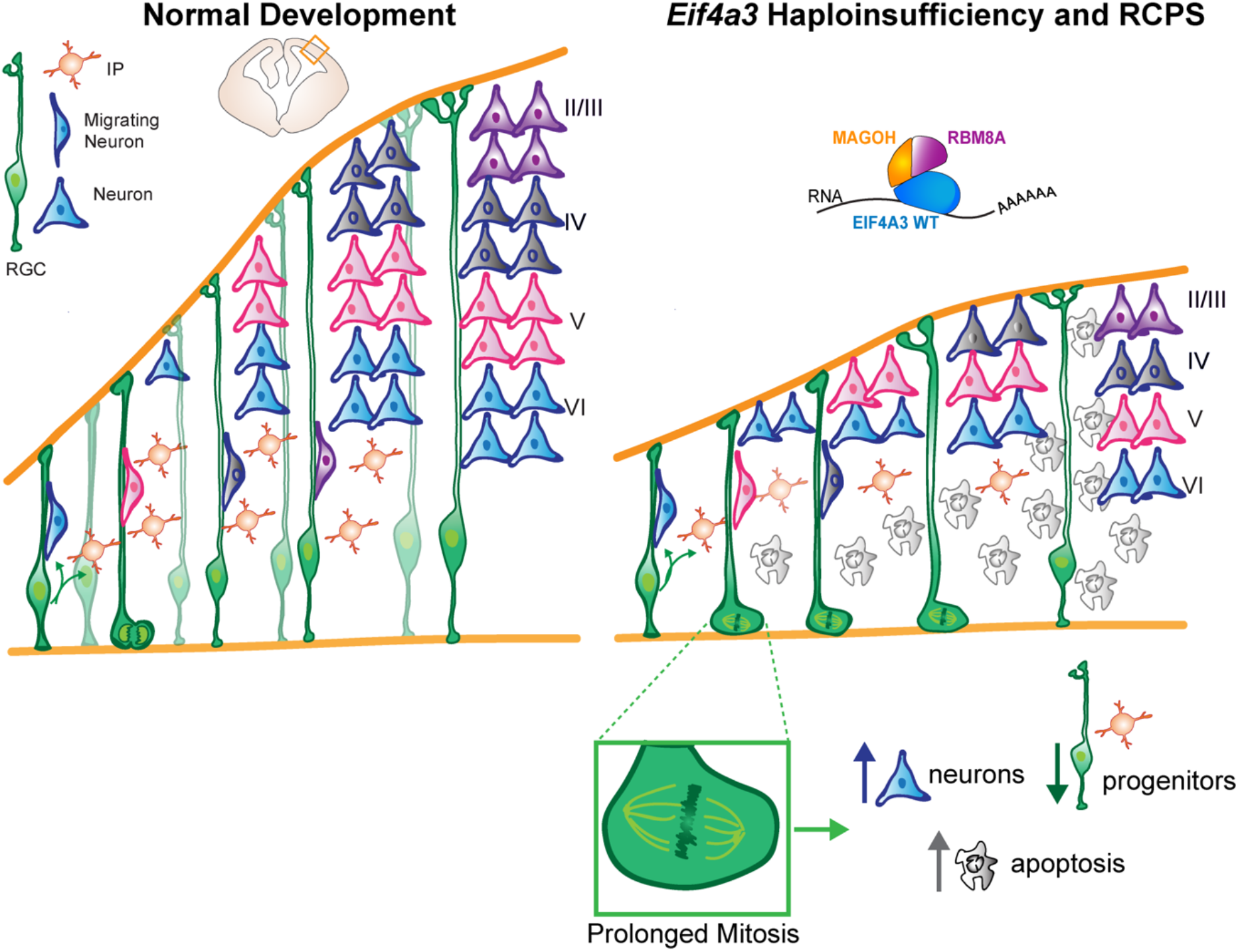
Cartoon model for *Eif4a3* control of neurogenesis. Graphical summary depicting normal development (left) and aberrant neurogenesis in a *Eif4a3* haploinsufficient mouse or RPCS model (right). *Eif4a3* haploinsufficient progenitors delayed in mitosis produce fewer progenitors, more neurons, and more apoptotic progeny. Our data suggest EIF4A3 controls neuron neurogenesis through both p53-dependent and independent mechanisms which diverge over the course of development.

## Discussion

The exon junction complex is essential for virtually every step of RNA metabolism across eukaryotes. Moreover, EJC components are critical for cortical development and underlie neurological disease. Using mouse models and human cortical organoids, we define key cellular and molecular mechanisms by which the RNA helicase component, EIF4A3, functions in cortical development and disease. EIF4A3 controls mitosis duration of both mouse and human progenitors to influence neuronal generation and survival. We further parse out developmental mechanisms of neurogenesis which are both *p53*-dependent and independent. Finally, we show that control of neuronal generation by EIF4A3 relies upon an intact EJC. Taken together, our study furthers an understanding of how EIF4A3 and the EJC control cortical development and suggests possible mechanisms by which they cause neurodevelopmental disorders.

### EIF4A3 controls mitosis duration to influence cell fate

We demonstrate that *Eif4a3* haploinsufficiency in mouse and human neural progenitors impairs neuronal generation and cell survival, and, strikingly, that these defects are associated with a prolonged mitosis. *Eif4a3*-depleted progenitors delayed in mitosis undergo fewer successful divisions and produce fewer progenitors, more neurons, and less viable progeny. These data corroborate previous findings from our group showing that mitotically delayed *Magoh* haploinsufficient RGCs or interneuron progenitors also exhibit altered neurogenesis (Pilaz et al., 2016; Sheehan et al., 2020). Our previous experiments support a causal role for mitosis in cell fate and microcephaly. Indeed using pharmacology to transiently prolong mitosis *in vitro, ex vivo* or *in vivo*, we demonstrated prolonged mitosis of mouse neural progenitors directly alters progeny cell fate (Mitchell-Dick et al., 2019; Pilaz et al., 2016). We have also shown a causal relationship for human neural progenitors (Mitchell-Dick et al., 2019). Thus, the current data for *Eif4a3* reinforce the relevance of mitosis duration for neural fate decisions.

A key question is how EIF4A3 controls mitosis duration. One possibility is that EIF4A3, through the EJC, post-transcriptionally controls genes necessary for mitosis progression. For example, in mouse embryonic stem cells (ESCs), EIF4A3 post-transcriptionally controls cell cycle regulators important for pluripotency maintenance (Li et al., 2022). Additionally, EIF4A3 directly binds transcripts encoding the cell cycle regulator *Ccnb1* to mediate its nuclear export in ESCs. In support of EJC-dependent post-transcriptional control of mitosis, all three core components are essential for mitosis (Pilaz et al., 2016; Sheehan et al., 2020; Silver et al., 2010). A second intriguing possibility is that EIF4A3 directly influences mitosis by manipulating the cytoskeleton. Indeed, EIF4A3 can directly bind to microtubules and also localizes to the mitotic spindle (Alsina et al., 2022). Likewise, RBM8A and MAGOH localize to centrosomes (Ishigaki et al., 2014; O’Neill et al., 2022). Thus it is possible the EJC could influence mitosis duration by directly modulating mitotic spindle integrity (Silver et al., 2010). Future detailed studies will be invaluable for teasing apart these potential canonical and non-canonical mechanisms.

More broadly, our findings support the notion that prolonged mitosis of progenitors is a causal mechanism for some microcephaly cases. To date, more than 25 genes encoding mitotic regulators have been implicated in primary microcephaly (Degrassi et al., 2019). Mitotic defects are present in many experimental microcephaly models, including immortalized cells, organoids, and mice (Jayaraman et al., 2018; Phan and Holland, 2021). For example, a study of Miller-Dieker syndrome (MDS) used patient iPSC-derived cortical organoids to show that delayed mitosis of outer radial glia is associated with aberrant neurogenesis (Bershteyn et al., 2017). Future studies will be invaluable to understand how mitosis duration influences fate and viability to shape development.

### Disentangling apoptosis and cell fate in *EIF4A3-*mediated microcephaly

*Eif4a3* haploinsufficiency in progenitors leads to striking microcephaly, associated with altered cell composition and extensive apoptosis. Thus, a key question is the extent to which these cell composition changes and microcephaly are explained by apoptosis. By compound ablation of *p53*, we show that microcephaly is due to both p53-dependent apoptosis and p53-independent mechanisms. We discover that *p53* ablation is sufficient to rescue deep-layer (Layers V-VI), but not upper-layer (IV-II/III) neurons. This finding is consistent with previous analyses of E18.5 *Rbm8a;p53* compound mutant brains which showed that *p53* loss rescued layer VI but not layer II/III neuron number (Mao et al., 2016). Probing E13.5, we discover that ablation of *p53* significantly rescues deep-layer generating progenitors, but is insufficient to fully rescue upper-layer generating progenitors at E16.5. These developmental timing differences suggest that progenitors themselves may have differential sensitivity to apoptosis over the course of development. It further suggests that EIF4A3-mediated control of neurogenesis relies on several downstream pathways, with p53 signaling being especially important for deep-layer neurogenesis, but additional yet undiscovered pathways relevant for fate of upper-layer neurons. Interestingly our p53 findings resemble that of some microcephaly mutants in which p53 partially but not completely rescues brain size (Little et al., 2021; Shi et al., 2019), whereas other mutants do show a complete rescue by this pathway (Insolera et al., 2014). This further highlights p53 as a key downstream pathway involved in microcephaly. Future studies, including transcriptomics, epigenomics, and proteomics, will be valuable to tease out how EIF4A3 differentially controls neurogenesis of upper- and deep-layer neurons. Further, while we have shown that mitosis is a key feature affected by *Eif4a3* mutation, the extent to which p53 signaling acts upstream or downstream of mitosis is a question for future studies.

### Unique requirements for EIF4A3 across cell types, developmental time, and disease

As a component of the EJC, EIF4A3 canonically controls RNA metabolism. Our rescue experiments argue that EIF4A3 acts in an EJC-dependent manner to control neuron number. This fits with the observation that *Eif4a3, Rbm8a*, and *Magoh* haploinsufficiency each cause similar cortical development phenotypes, including microcephaly, precocious neurogenesis, apoptosis, and mitotic defects. Further, all 3 mutants exhibit converging dysregulation of common transcripts (Mao et al., 2016). Thus, we postulate that for neurogenesis, EJC components work together. In contrast, recent findings from our lab indicate there are EJC-independent requirements for EIF4A3 in neurons; *Eif4a3*, but not *Magoh* or *Rbm8a*, is required for axonal outgrowth (Alsina et al., 2022). We find that EIF4A3 is competent to bind to microtubules independent of the EJC and RNA, and directly promotes microtubule polymerization and stability (Alsina et al., 2022). This raises the intriguing possibility that EIF4A3 could also have some EJC-independent roles in neurogenesis, such as at later stages of development not examined in this study.

Although *Eif4a3* has conserved roles in the production of excitatory neurons in both mouse and human neural progenitors, it is intriguing to consider if this holds true in other brain regions. In both excitatory and inhibitory progenitors, *Magoh* is required for proper mitotic progression and neurogenesis, illustrating parallel functions at play in the cortex and non-cortical regions (Pilaz et al., 2016; Sheehan et al., 2020). In future studies it will be interesting to assess if all EJC components similarly control inhibitory neurogenesis and whether they do so via canonical mechanisms.

Mutations in *EIF4A3* and *RBM8A* are linked to neurodevelopmental disorders including microcephaly, intellectual disability, and RCPS (Bertola et al., 2018; Favaro et al., 2014; Hsia et al., 2018; Nguyen et al., 2013). Our analysis of cortical organoids and dissociated neural progenitors reveal that neurogenesis is a key process impacted in RCPS. In particular, we note conserved roles for *EIF4A3* in mitotic duration, progeny fate, and apoptosis. Our findings complement previous observations that axonal maturation is impaired in RCPS cortical organoids (Alsina et al., 2022). Continued investigation of RCPS organoids can give important insights into the etiology of this rare disorder. In sum, we propose a model in which *Eif4a3* haploinsufficiency causes microcephaly both through apoptosis and altered cell fate due to mitotic delay. Altogether, our study provides new insights into the basis for RCPS and other EJC-dependent disorders.

## Acknowledgements

We thank the members of the Silver Lab for helpful discussions and careful reading of the manuscript. We thank the Duke Mouse Transgenic Facility. We are grateful to Maria Rita Passos-Bueno and Gershon Kobayashi for the RCPS iPSCs.

## Author contributions

BML and DLS conceived of the project and wrote the paper. DLS oversaw research. BML and RAS performed analyses of *Eif4a3* mutant brains. BML performed live imaging experiments. CMM generated cortical organoids with the help of BML. BML performed cortical organoid experiments. BML performed rescue experiments, with assistance from FCA.

## Materials & Methods

### Mouse husbandry and genetics

All animal procedures were approved by the Duke Institutional Animal Care and Use Committee (IACUC) and performed in agreement with the ethical guidelines of the Division of Laboratory Animal Resources (DLAR) from Duke University. We used the previously described mouse lines: *Eif4a3*^loxP^ and *Emx1-Cre (B6*.*129S2-Emx1tm1(cre)Krj/J)* (Gorski *et al*, 2002). The following mouse strains were obtained from Jackson Laboratories: C57BL/6J (*wild type*) and B6.129S2-Trp53^tm1Tyj^/J (p53^LoxP^). For embryo staging, plug dates were defined as embryonic day (E) 0.5 on the morning the plug was identified.

### Immunofluorescence

#### Embryonic brains

Embryonic brains were fixed and sectioned as previously (Mao et al., 2015). Coronal 20 μm sections from the somatosensory cortex were permeabilized with 1X PBS/0.25% TritonX-100 and blocked with 5% NGS/PBS for 1 hour at room temperature. Sections were incubated with primary antibodies overnight at 4°C, and secondary antibodies at room temperature for 2 hours (Alexa Fluor-conjugated, Thermo Fisher, 1:500). The following primary antibodies were used: SOX2 (Thermo Fisher, 14-9811-82, 1:1000), CTIP2 (Abcam, c8035, 1:500), TBR2 (Abcam, AB23345, 1:1000), CC3 (Cell Signaling, 9661, 1:250), PH3 (Millipore, 06–570, 1:500), TBR1 (Cell Signaling Technology, 49,661 S, 1:1000), LHX2 (Millipore, ABE1402, 1:500). Slides were mounted with Vectashield (Vector Labs, H-1000–10). For antigen retrieval of ROR (R&D systems, N7927, Lot #A-2, 1:100), coronal 20 μm sections were boiled for 15 minutes in sodium citrate buffer (10mM sodium citrate, 0.05% Tween-20, pH 6.0) in a histology slide container. Slides were taken off the heat but kept in sodium citrate buffer an additional 10 minutes to cool. Slides were then removed from the container and allowed to cool an additional 10 minutes before being rinsed 3 times in 1X PBS. Permeabilization and antibody incubations were then followed as described above.

#### Cortical organoids

Organoids were fixed in cold 4% PFA/PBS for 2 hours, rinsed 3 times in cold 1X PBS, and submerged in 30% sucrose/PBS overnight. Organoids were embedded, sectioned, and stained as described above for embryonic brain tissue. SOX2, PH3 antibodies were used as described above, as well as Ki67 (Cell Signaling Technology, 12202, 1:250).

#### Imaging and analysis

Images were captured using a Zeiss Axio Observer Z.1 equipped with an Apotome for optical sectioning, at 5X, 10X, and/or 20X. Between two and three sections were imaged per embryo or organoid, and all images for a given experiment were captured with identical exposures. Images were cropped to 300 μm radial columns and the brightness was equivalently adjusted across all images in Fiji. Cells were either manually (Fiji cell counter) or automatically (QuPath) counted. QuPath parameters were used as previously (Hoye et al., 2022). For binning analysis, 300 μm wide radial columns were divided into 5 evenly spaced bins spanning from the ventricular (bin 1) to the pial (bin 5) surface as previously (Hoye et al., 2022). Each cell was assigned to a bin to calculate the distribution.

#### Primary cultures and live imaging

Primary cortical cultures were derived from E12.5 embryonic dorsal cortices, as previously described (Mitchell-Dick et al., 2019), but with minor modifications: 1) cortices were trypsinized for 5 minutes and 2) 150,000 cells were plated on poly-D-lysine coated glass-bottom 24-well culture plates (MatTek). Images were captured every 10 minutes as previously described (Pilaz et al., 2016). Mitosis duration and cell division were identified by morphology (rounding and condensation of chromatin). Fate determination was performed post-imaging by immunostaining for Tuj1, Sox2, and Tbr2, as previously described (Mitchell-Dick et al., 2019). Live imaging of dissociated human progenitors was performed similarly, except fate determination was performed post-imaging by immunostaining for anti-Ki67 (Cell Signaling Technology, 12202, 1:1000) and Tuj1 (TUBB3, Biolegend, 801202, 1:2000).

#### Generation of brain cortical organoids

Work with H9 cells and iPSCs was approved by the Duke University Institutional Research Board. The H9 (WA09) human ES cell line was purchased from WiCell (hPSCReg ID: WAe009-A) (Thomson et al., 1998). The control and RCPS patient iPSC lines used in this study were previously characterized and genotyped (Alsina et al., 2022; Miller et al., 2017; Yoon et al., 2019) and the authors are blind to patient identity. The isogenic iPSC line was previously generated and characterized (Alsina et al., 2022). Cells were maintained in Essential 8 medium (Gibco) supplemented with 100 μg/ml Normocin (Invivogen) on Matrigel (Corning)-coated plates, and then seeded on vitronectin (Gibco) for 2 passages before generating the 3D cultures. Cortical organoids were generated and maintained as previously described (Alsina et al., 2022; Yoon et al., 2019). Images were taken at 4X magnification using the EVOS XL Core (ThermoFisher) microscope (scale: 2.145 pixels/μm).

#### Dissociation and electroporation of organoids

Dissociation of H9 organoids into 2D cultures was performed as previously described with minor modifications (Miura et al., 2020). To electroporate the single cell suspension, the standard protocol for the Lonza P3 primary cell 4D Nucleofector X Kit S was followed using pulse code CB-150. For the electroporation, 1 × 10^6^ cells and 1.5uL of 10 μM siRNA mix were used for each well in the 16-well cuvette. The following siRNAs were used from QIAGEN: Scrambled siRNA, Hs_EIF4A3_2 (Cat No. SI00107828), Hs_EIF4A3_3 (Cat No. SI00107835), Hs_EIF4A3_5 (Cat No. SI02663794) and Hs_EIF4A3_6 (Cat No. SI03049676). 25nM of each *EIF4A3* siRNA was mixed together for a total of 100nM stock solution. Stock was diluted 1:100 to 10 μM before electroporation. 5 × 10^4^ cells were seeded per well in 24-well MatTek glass-bottom plates (P24G-1.5-10-F) coated with 0.1 mg/ml poly-D-lysine (Sigma; 30,000-70,000 mol wt). Cultures were maintained in organoid base medium without small molecules. After 48 hours of knockdown, live imaging was started. Knockdown of *EIF4A3* was confirmed by RT-qPCR (Figure 5B).

#### EdU in organoids

Organoids were incubated in media containing 10mM EdU for 48 hours. Organoids were fixed as described previously (Mitchell-Dick et al., 2019). Click-It reaction was performed using Life Technologies Click-It Plus Kit (#C10749) following the manufacturer’s instructions. Antibody staining was then performed as described above.

#### Electroporation of primary progenitor cultures

Primary cortical cultures were derived from E12.5 embryonic dorsal cortices as described above. The standard protocol for the Lonza P3 primary cell 4D Nucleofector X Kit S was followed using pulse code CM-150 and 0.5 × 10^6^ cells per electroporation in the 16-well cuvette. Three different electroporation conditions were used per sample: 1μg pCAG-GFP only, 1 μg pCAG-GFP + 0.25ug pCAG-3xFLAG-Eif4a3 (*Mus musculus*), and 1 μg pCAG-GFP + 0.25 μg pCAG-3xFLAG-Eif4a3 T163D (*Mus musculus*) (Alsina et al., 2022). Concentrations of pCAG constructs were empirically determined to avoid overexpression phenotypes. The primary cultures were incubated for 48 hours before fixation and immunostaining for Tuj1 as previously.

#### RT-qPCR and primers

RNA was extracted in RLT buffer plus 0.01% β-mercaptoethanol following RNeasy kit (QIAGEN) and cDNA was prepared using the iScript kit (Bio-Rad) following manufacturer’s protocol. qPCR was performed using SYBR Green iTaq (Bio-Rad) in at least 3 independent biological samples (3 technical replicates per sample) in a QuantStudio 3 machine (Applied Biosystem). Values were normalized to *TBP* as loading control. The following primers were used: *Homo sapiens TBP* (Forward 5’-GTGACCCAGCATCACTGTTTC-3’ and Reverse 5’-GCAAACCAGAAACCCTTGCG-3’) and *Homo sapiens EIF4A3* (Forward 5’-GGAGATCAGGTCGATACGGC-3’ and Reverse 5’-GATCAGCAACGTTCATCGGC-3’).

## Statistical methods and rigor

Exact statistical tests, p-values, and n for each analysis are reported in **Supplemental Excel 1**. For each experiment, both male and female mice were used and littermates were used when possible. All analyses were performed by 1 or more blinded investigators.

**Supplemental Figure 1.**
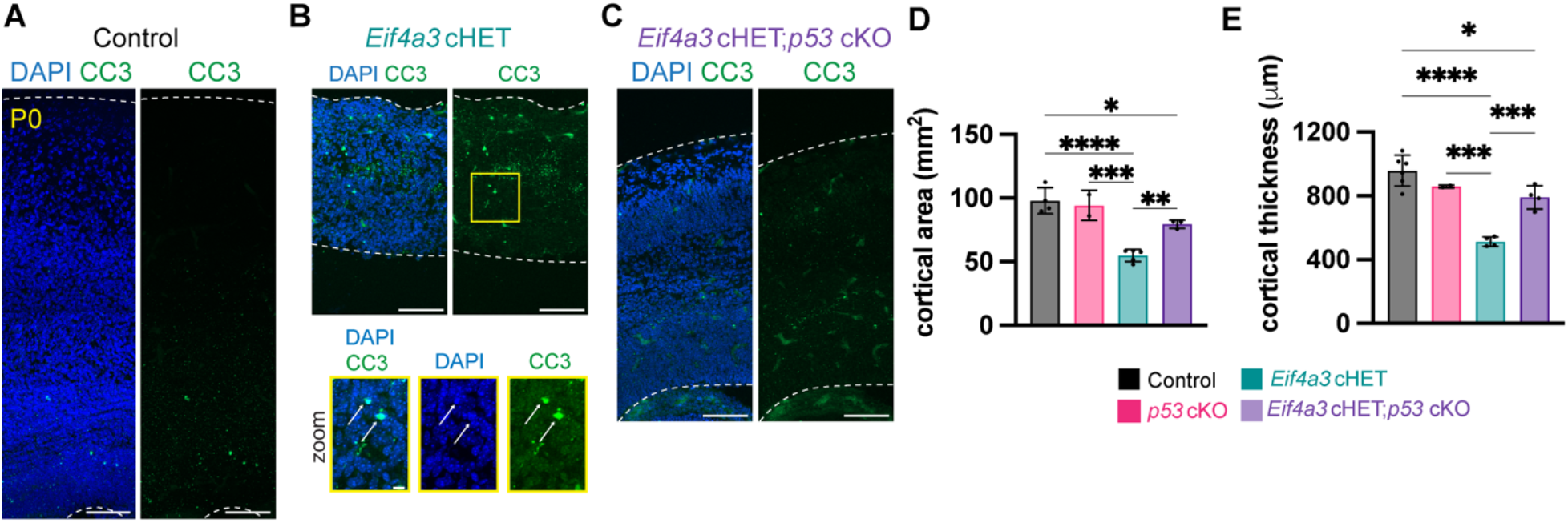
Analysis of *p53;Eif4a3* compound mutant at P0. **(A-C)** Representative sections of P0 cortices for indicated genotypes stained with CC3 (green) and DAPI (blue). **(B)** Zoom panel of *Eif4a3* cHET sections stained with CC3 (green) and DAPI (blue). Arrows indicate CC3+ and DAPI+ double-positive cells. (**D)** Quantification of cortical area and **(E)** cortical thickness in P0 brains with indicated genotypes. N=3-7 embryos per genotype for **(A-E)**. Dotted lines demarcate dorsal cortex. ANOVA with Tukey post-hoc: *P<0.05, **P<0.01, ***P<0.00, ****P<0.001, ns=not significant. Individual dots represent biological replicates and error bars represent s.d. Scale bars: 100 μm (**A-C)**, 10 μm (**B zoom**).

**Supplemental Figure 2.**
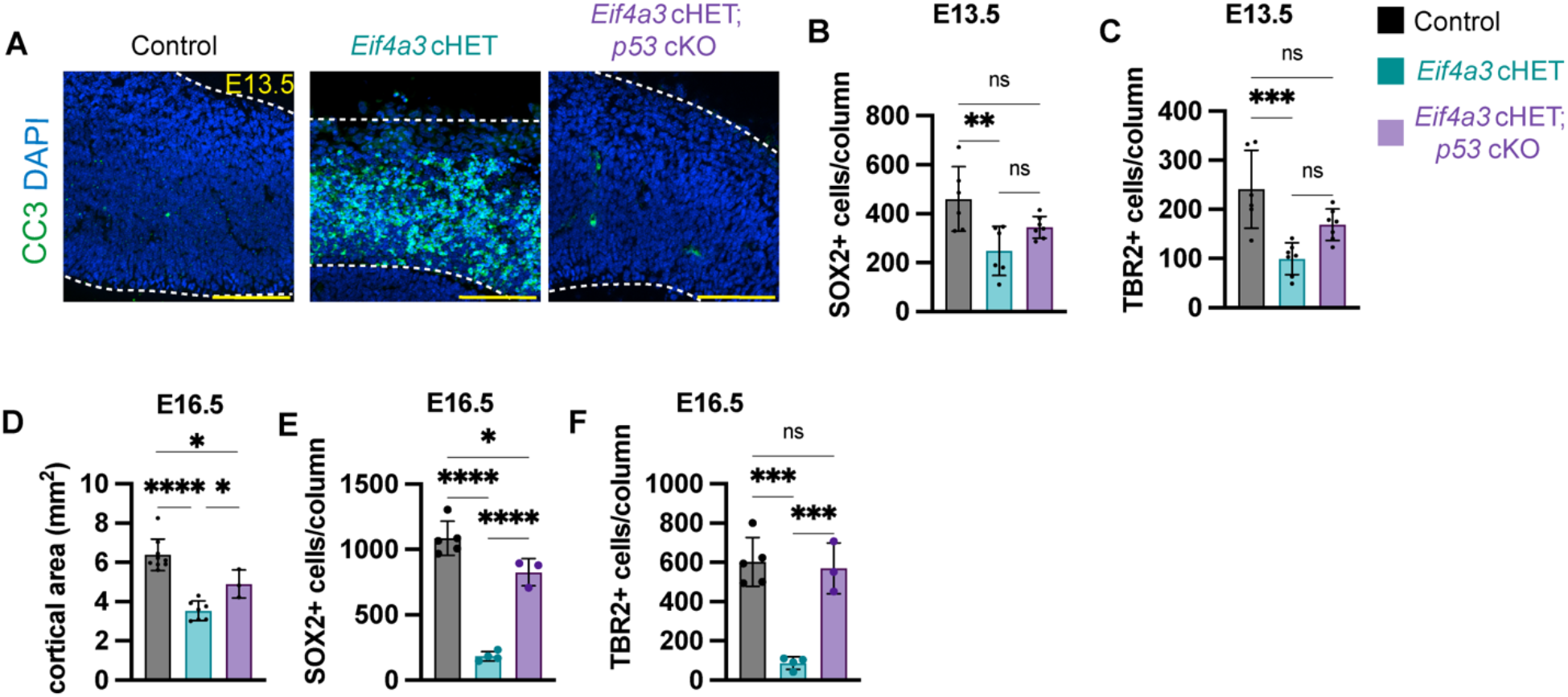
Analysis of *p53;Eif4a3* compound mutant at E13.5 and E16.5. **(A)** Representative sections at E13.5 of indicated genotypes stained with CC3 (green) and DAPI (blue). **(B,C)** Quantification of total number of **(B)** SOX2+ cells **(Figure 5C)** and **(C)** TBR2+ cells **(Figure 5D)** in 300 μm cortical columns at E13.5. **(D)** Quantification of cortical area in E16.5 brains with indicated genotypes. **(E,F)** Quantification of total number of **(E)** SOX2+ cells **(Figure 5I)** and **(F)** TBR2+ cells **(Figure 5K)** in 300 μm cortical columns at E16.5. For E13.5: N=7-10 embryos per genotype for **(A-C)**. For E16.5: N=3-5 embryos per genotype for **(D-F)**. Dotted lines demarcate dorsal cortex. ANOVA with Tukey post-hoc: *P<0.05, **P<0.01, ***P<0.001, ****P<0.001, ns=not significant. Individual dots represent biological replicates and error bars represent s.d. Scale bars: 100 μm **(A)**.

**Supplemental Figure 3.**
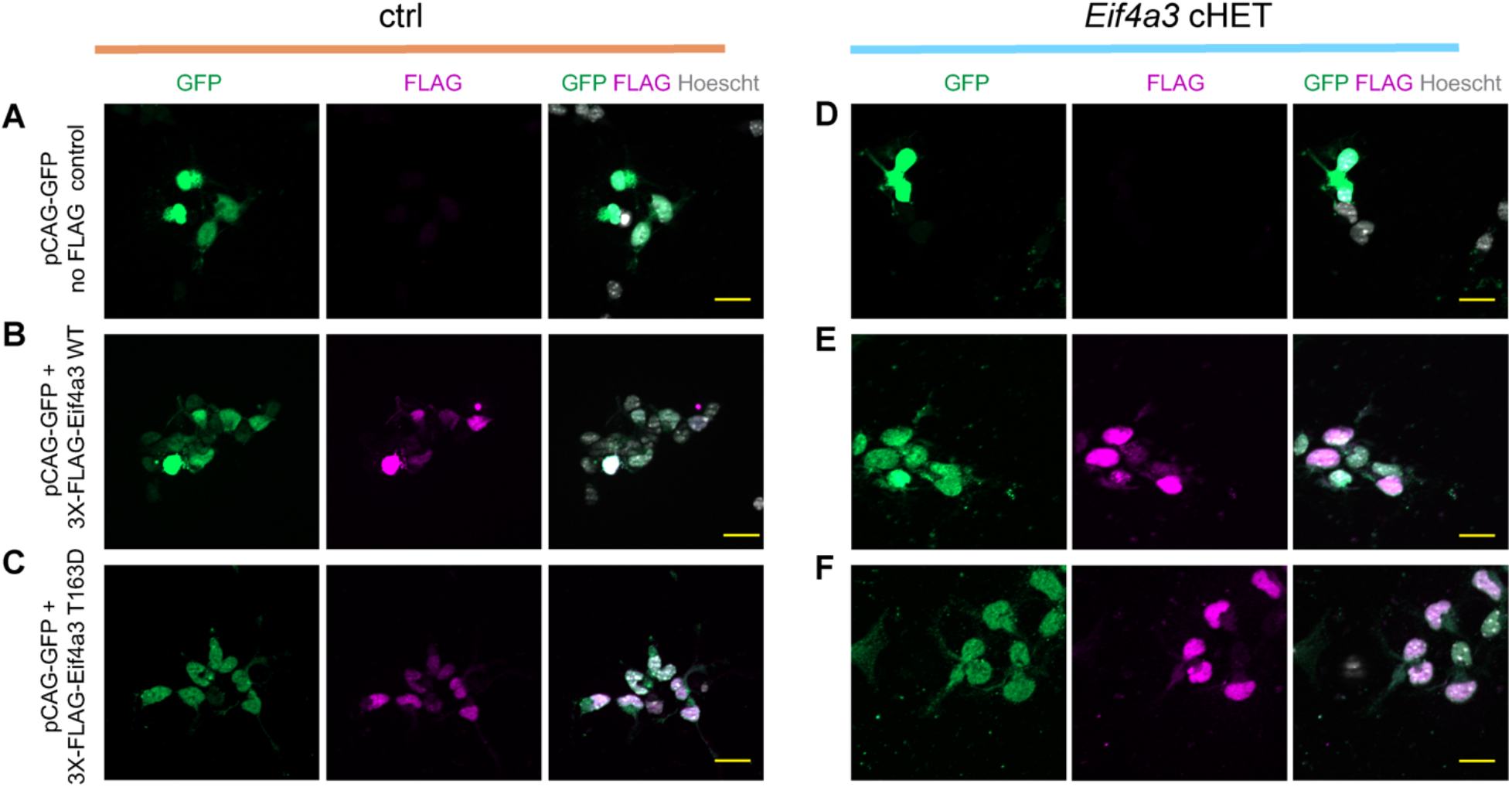
WT and T163D mutant EIF4A3 constructs are expressed in control and cHET E12.5 progenitor cultures. **(A-C)** E12.5 control (*Emx1*-Cre;*Eif4a3*^+/+^) or **(D-F)** cHET (*Emx1*-Cre;*Eif4a3*^*l*ox/+^) primary cultures were electroporated with pCAG-GFP and the pCAG-3x-FLAG constructs depicted in **Figure 8A**. To test for expression of the plasmids, cultures were fixed and stained for GFP (green), FLAG (magenta), and Hoescht (white). Left panels show GFP and Hoescht, middle panels show FLAG and Hoescht, and right panels show the merge of GFP, FLAG, and Hoescht. Scale bars: 20 μm.

